# Body size and cranial shape differentiation in urban and rural house mice (*Mus musculus domesticus*)

**DOI:** 10.64898/2026.05.16.725634

**Authors:** Stephen C. Kupchella, Anne E. Kort, Megan Phifer-Rixey

## Abstract

Cities are characterized by elevated temperatures, increased pollution, and high-density human populations which often are accompanied by changes in available resources, like food. These shifts have the potential to drive phenotypic divergence in urban wildlife. Functional morphological traits, like body size, can mediate interactions between wildlife and habitat and are closely tied to life history and fitness. While examples of functional morphological variation associated with urbanization are increasing, variation in such traits as a response to urbanization remains unexplored for most taxa. Here, we investigated morphological divergence between urban and rural populations of house mice (*Mus musculus domesticus*). House mice are globally distributed in diverse habitats and are a model system with a wealth of phenotypic data, making them useful for the study of the impacts of urbanization on morphology. Using a paired replicate design, we sampled urban and rural populations in three distinct metropolitan regions in the eastern United States. We found that body size was smaller in urban populations. Using 3D geometric morphometrics, we also analyzed variation in cranial shape across habitats. Differences in cranial shape were largely allometric, that is, driven by differences in body size. However, we also uncovered evidence of cranial shape variation between habitats not explained by size. In contrast, we did not find evidence for habitat-driven differences in cranial capacity independent of size. Overall, our results suggest a key role for body size in mediating morphological responses to urbanization and highlight the potential of house mice as a globally-distributed model for urbanization.

## 1. INTRODUCTION

Urbanization represents a significant and ongoing transformation of natural ecosystems. Globally, urban areas are growing rapidly, with estimates ranging between 0.6 – 1.3 million km^2^ by the year 2050 (Huang et al. 2019). This expansion has altered the structure and function of landscapes and produced novel habitats that differ markedly from the natural areas they replace. Wildlife that persist in cities are subject to a suite of anthropogenic stressors including pollution (chemical, sound, light), elevated temperatures, and altered resources like food and trash, among others. These factors can impact ecological dynamics, changing the ways animals interact with the environment and resulting in phenotypic divergence (Alberti et al. 2017; Thompson et al. 2022). Morphological traits are at the interface between organisms and the environment and thus can shed light on the impacts of urbanization on wildlife. Many such traits (e.g., body size, cranial and appendage shape) are closely tied to performance and fitness, and divergence in morphological traits across habitats can signal shifts in function, pointing to potential eco-evolutionary drivers. For example, urban house finches (*Haemorhous mexicanus*) have longer and narrower beaks than rural conspecifics. Evidence suggests that individuals with longer bills can sing at higher frequencies, overcoming the degradation of acoustic signals caused by low-frequency noise pollution. Additionally, beak shape may be influenced by feeding ecology, with the urban phenotype better suited for the manipulation of large seeds common to backyard bird feeders in urban areas (Giraudeau et al. 2014).

Examples of intraspecific morphological differentiation associated with urbanization are growing, with data for a range of vertebrate taxa including birds, reptiles, and mammals (Giraudeau et al. 2014; Winchell et al. 2016; Yu et al. 2017; DePasquale et al. 2020; Falvey et al. 2020; Gangloff et al. 2025; see also Johnson and Munshi-South 2017). In particular, studies have investigated variation in body size. Body size has the potential to influence multiple aspects of a species’ ecology and life history, including thermoregulation, metabolic rate, home range size, predator-prey dynamics, competition, fecundity, and resource acquisition (Kleiber 1932; Blueweiss et al. 1978; Western 1979; Cohen et al. 1993). Within mammals, rodents are the most diverse order and have become important for understanding the impacts of urbanization on biodiversity, as both native and non-native commensal species occupy urban habitats (Castillo et al. 2003; Cavia et al. 2009; Garba et al. 2014; Larson and Sander 2025). Body size differentiation associated with urbanization has now been documented in multiple rodent systems. However, there is no clear consensus on whether cities are associated with larger or smaller body size in rodents. Some studies have found that animals are smaller bodied in urban areas (Wist et al. 2022; Bateman et al. 2023; Ofori et al. 2024). Such shifts would be consistent with predictions of Bergmann’s rule in the context of urban heat islands (UHI), but also with other potential ecological drivers, e.g. response to predation. Other studies have documented size increases (Pergams and Lawler 2009; Pieniążek et al. 2017; Tranquillo et al. 2024) and highlight potential drivers such as resource availability and dispersal ability (Martin and Sheridan 2022). In addition, studies vary widely approach ranging studies of single taxa in a single city to metanalyses of broad taxonomic groups.

Craniofacial variation associated with urbanization has also been shown in several rodent species (Pergams and Lacy 2008; Pergams and Lawler 2009; Snell-Rood and Wick 2013; Yu et al. 2017; DePasquale et al. 2020; Puckett et al. 2020; Feijó et al. 2025). For example, some studies have reported a trend toward increased cranial capacity in urban environments, purportedly induced by plastic and/or adaptive responses to structural complexity (Snell-Rood and Wick 2013; DePasquale et al. 2020). However, evidence supports the opposite trend in some taxa (Snell-Rood and Wick 2013), and other studies have failed to detect a signal altogether (Puckett et al. 2020). Aside from the brain, the cranial skeleton houses organs used in sensory transduction (e.g. vision, olfaction, audition) as well as structures critical for resource acquisition and mastication. Dietary ecology is associated with biomechanical changes in loading and bite force capacity that can be detected in the shape of the skull (Maestri et al. 2016; Verde Arregoitia et al. 2017). As such, shape variation in the skull between habitats may indicate potential differences in function, particularly with respect to diet.

House mice (*Mus musculus domesticus*) have great potential as a model for studying morphological responses to urbanization. Wild house mice are ubiquitous in urban areas, have short generation times, and their use as a laboratory model has produced an abundance of phenotypic and genetic resources. Here, we investigated morphological differentiation between urban and rural populations of house mice in three metro regions in the eastern United States (New York, NY., Philadelphia, PA., and Richmond, VA). Previous research has shown that mice from these areas are genetically distinct (Phifer-Rixey et al. 2018). Therefore, this paired replicate design allowed us to identify differences among habitats in each region and evaluate to what extent differences are shared among regions, potentially reflecting parallel responses to urbanization (Johnson and Munshi-South 2017; Rivkin et al. 2019). First, we tested for differences in body size, examining both weight and linear size metrics, finding that urban mice are smaller than rural mice. Next, we used 3D geometric morphometrics to quantify and compare the shape of the cranium between habitats. We found significant differences in cranial shape across habitat. Differences were largely allometric, but we also found evidence of cranial shape variation between habitats independent of size. Finally, we tested whether urban and rural populations of house mice differ in their cranial capacity, finding no evidence of differences in cranial capacity beyond those expected due to size. Overall, our results suggest that urban and rural mice differ significantly in body size, a trait with strong ties to fitness via physiology, metabolism, and ecological interactions, and correlated traits like skull size and shape which also have potential to impact fitness.

## 2. Materials and Methods

### [2.1] Sampling

House mice were collected using live traps (H.B. Sherman, FL) from three cities in the eastern United States (NYC: New York, NY., PHL: Philadelphia, PA., RVA: Richmond, VA.; FIG 1; S1 Table) and the rural areas surrounding them. Trapping areas were defined as urban or rural based on the Global Human Settlement Layer (GHS-SMOD) settlement classification (Schiavina et al. 2023; Pesaresi et al. 2024). Upon collection, mice were euthanized and processed with the approval of the Drexel University Institutional Animal Care and Use Committee (IACUC; Protocol No. LL-122-74) and the Monmouth University IACUC (Protocol No. Asp1802). Standard phenotypic measurements (weight, length, hind-foot length, and tail length) were collected for most specimens. Weight was measured using a digital scale (Ohaus Scout SPX421, ± 0.001g), and length measurements were recorded with a ruler (± 1mm). For a subset of specimens, skins, skeletons, and skulls were prepared and deposited at the Yale Peabody Museum in New Haven, CT (S1 Table). After processing, we used digital calipers to obtain length measurements of the left femur from the superior surface of the femoral head to the inferior surface of the medial condyle (± 0.01mm). The mediolateral width of each femoral shaft was also collected at the midpoint of the diaphysis. For specimens with broken left femora, we recorded contralateral measurements on the right femur. Femoral length measurements were recorded three times, and the average of the three was retained for analysis.

**Figure 1.**
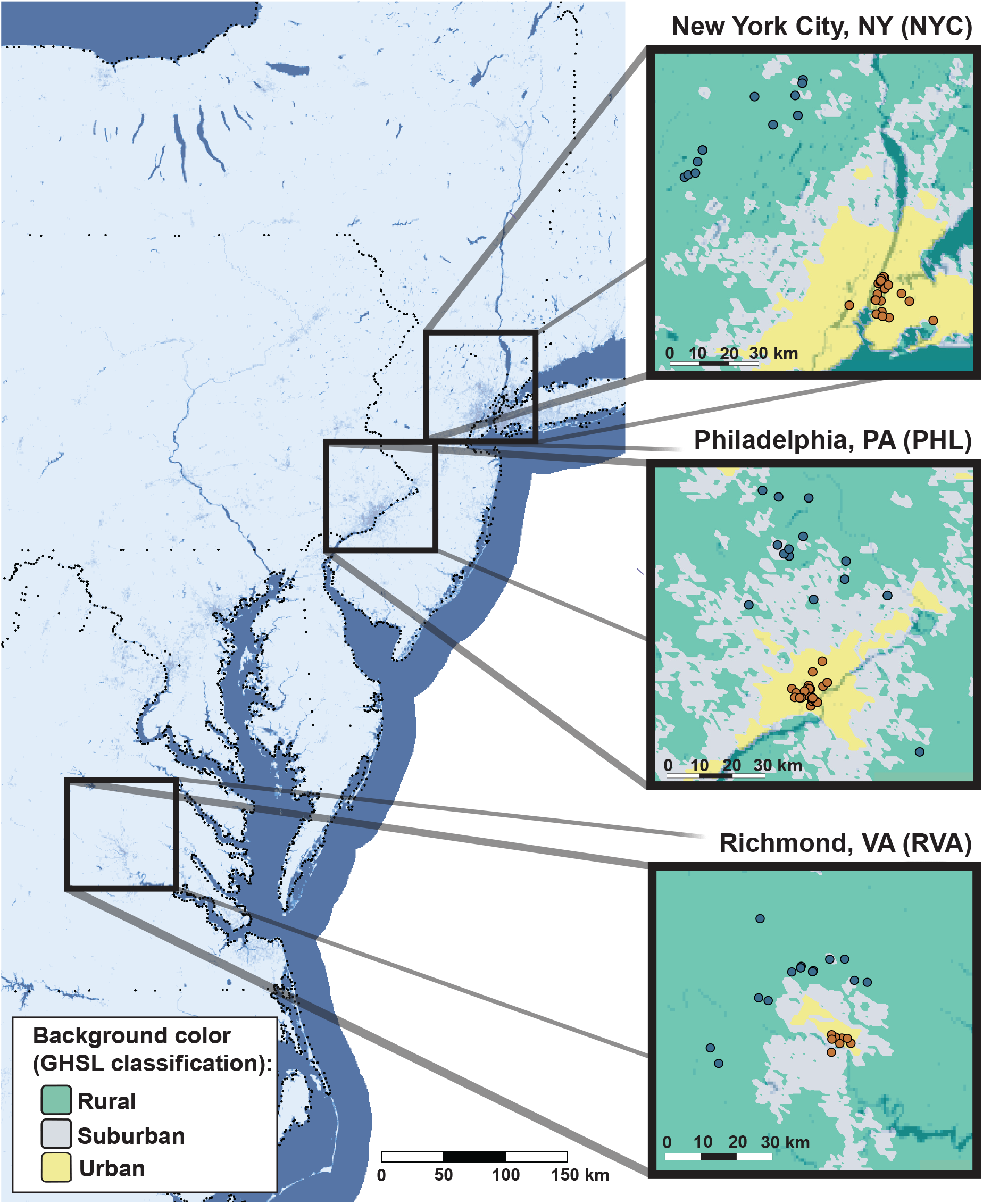
Map of sampling sites. House mice were collected from highly developed and less developed sites in three metro regions: New York City, NY., Philadelphia, PA., and Richmond, VA. Inset background color indicates degree of urbanization (GHSL-SMOD (Pesaresi et al., 2024; Schiavina et al., 2023)). Within insets, orange points indicate urban sites and blue points indicate rural sites. Figure adapted from Giancarli et al. 2026.

Mice used in this analysis were collected from June 2022 to March 2025. To minimize the likelihood of including related individuals in the datasets, we prioritized collection from structures and properties that were >500m apart. If more than one mouse was collected from the same location, only one was retained in the analysis with the decision based on balancing the collection of data for both sexes. It is not possible to obtain accurate age estimates for wild-caught specimens. As a result, we used a general developmental indicator (hindfoot length ≥18mm) to categorize samples as adult, removing all individuals with lower values with three exceptions. Two specimens had no recorded hindfoot measurement but were otherwise typical for adults and thus were included. In addition, one specimen with a hindfoot measurement of 17mm was a retained based on body weight which clearly placed it as an adult (19.62g vs 15.71g (average); S1 Table). Additionally, multiple pregnant females included in the initial collection were omitted from body weight analyses. Because of these considerations, and because some traits could not be collected for all individuals due to damage, we created three distinct datasets for each of the major morphological analyses: weight (n = 80), linear size traits (n = 81), and cranial traits (n = 72).

### [2.2] Analysis of body mass and linear size measurements

We analyzed multiple aspects of body size, including body mass, body condition, body length, femur length, and femoral mediolateral width by fitting GLMs in RStudio v 4.5.1 (R Core Team 2024). We investigated body mass, body condition, and body length metrics to determine overall size. Femoral length was chosen as a proxy for body length (Melin et al. 2005), with femoral mediolateral width serving as an indicator of potential differences in mechanical loading and/or locomotor ability (Wallace et al. 2012; Castro and Garland Jr. 2018). Model assumptions were diagnostically tested using the Shapiro-Wilk test for normality and Levene’s test for homogeneity of variance where appropriate. To test for significant relationships, Type II ANOVAs were implemented in the *car* package v. 3.1.3 (Fox and Weisberg 2019). All models were fit with three primary predictors: sex, habitat (urban/rural), and metro region (NYC, PHL, RVA). Where relevant, additional models were created with size metrics included as a covariate. For example, to evaluate the contribution of habitat to differences in weight, we included body length as a covariate to account for scaling. We further analyzed pairwise comparisons using post-hoc tests in the *emmeans* package to assess significant interaction terms and obtain estimated marginal means (EMMs) and differences in EMMs among groups.

### [2.3] Cranial Morphology

#### [2.3a] Micro-Computed Tomography

We used a Scanco Medical µCT 45 instrument (Scanco Medical, Brüttisellen, Switzerland) housed at the Penn Center for Musculoskeletal Disorders (University of Pennsylvania) to generate micro-CT image data for each available skull. Up to six mouse skulls were stacked 3×2 and packed with polyethylene foam into a 50ml conical vial to prevent movement during scanning. Specimens were scanned at 12µm resolution (55 kVp, 8W, 300ms). Using the scanning software, segments of the tube were isolated such that three skulls were scanned concurrently, resulting in two sets of image stacks per vial.

#### [2.3b] Landmarking

3D digital renderings of mouse crania were generated in *3D Slicer* (Kikinis et al. 2013). Of the mice collected, 95 adult crania were scanned. Of these, 16 with fractured or missing elements and seven from duplicate locations were excluded, leaving 72 skulls for comparative morphometric analysis. Each skull was digitized using an adapted fixed-landmark scheme originally published by Pallares et al. 2014. This scheme was selected for its broad anatomical coverage and potential to capture variation in overall cranial shape. Some changes to the scheme were made. One landmark that was difficult to reliably digitize was replaced with two points (RLP:LLP 2, 3) to capture variation in the width of the brain case. Additionally, two points on each zygomatic arch were dropped and replaced with 20-point semi-landmark curves to increase the density of points on a structure that lacks homologous landmarks (S2 Table, S1 Figure).

Due to the bilateral symmetry of the cranium, it is common to landmark one half of the structure and reflect points across the midsagittal plane. This approach was implemented to reduce placement error and limit asymmetric variance in the dataset. To achieve this, we first extracted and manually digitized landmarks that defined the midsagittal plane (MSP). We used a custom script executed in the 3D Slicer python terminal to create a plane of symmetry and reflect manually digitized right lateral points (RLP) and right zygomatic curves (RZC) across the plane, producing a mirrored left lateral point (LLP) and left zygomatic curve (LZC) point list (S2 Table, S1 Figure). The five-point lists were then merged using the Merge Markups utility in *Slicermorph* (Rolfe et al. 2021) to produce an 86-point data set capturing the complete shape configuration of each mouse skull. On a significant number of the skulls, one or both pterygoid processes were poorly rendered, reflecting incomplete capture during scanning. Thus, prior to testing, two landmarks originally digitized on the processes were removed, resulting in a final dataset of 84 landmarks. Supplementary Tables S1, S2, and Supplementary Figure S1 reflect the loss of these landmarks.

#### [2.3c] Endocasts

Endocranial segments of each skull were generated using the *Segment Endocranium* utility in Slicermorph with a smoothing kernel size of 2.00mm and maximum hole size of 4.00mm. The resulting cranial endocasts were visually inspected to ensure that regions rendered appropriately, and that no artifacts were present. If the first run of the software produced an endocast with irregular projections or artifacts emanating from the surface of the segment, the software was run again with the same parameters, and data were only retained if artifacts were resolved following the second iteration. Volumetric data was collected for each endocast in the *Segment Statistics* module.

#### [2.3d] Geometric Morphometrics

To test the effects of sex, habitat, and metro region on cranial variation, raw landmark coordinates were exported to RStudio. We first implemented a Generalized Procrustes Analysis (GPA) (Rohlf and Slice 1990), in the *geomorph* package v. 4.0.10 (Adams et al. 2012; Baken et al. 2021) using a generalized least squares criterion to remove rotation, translation, and scale. The GPA with partial Procrustes superimposition was performed on 48 fixed and 36 sliding semi-landmarks. The Procrustes aligned coordinates were then tested against the set of categorical variables with a Procrustes permutation ANOVA set to 1000 iterations using the ‘procD.lm’ function. The centroid size, calculated as the square root of the sum of squared distances of the landmark configurations from their geometric mean, was retained and added as a covariate to control for potential size and shape covariation.

We then extracted major axes of shape variation by performing a Principal Component Analysis (PCA) using the gr.prcomp function in *geomorph*. The highly dimensional nature of 3D coordinate data results in many defined principal components (PCs) that represent unstructured noise or fine-scale local variation. To identify PCs that correspond to major biological patterns of shape variation in the dataset, we performed a parallel analysis using Horn’s test of principal components with the *paran* package v. 1.5.4 (Dinno 2009). This technique compares observed eigenvalues to those generated from a randomly permuted null distribution, and only components whose eigenvalues exceeded the simulated threshold were retained for further analysis. We then used these PCs as response variables to test the effects of the predictors on shape while removing redundancy with MANCOVA.

#### [2.3e] Allometry

We tested and visualized two allometry models using the plotAllometry function to understand potential differences in the allometric slope and intercept between habitats. Models were fit with an interaction term (shape ∼ size * habitat) as well as an additive term (shape ∼ size + habitat) (S3 Figure). We then identified the common allometric vector and decomposed shape into allometric and non-allometric components using the ‘calculate common allometric component’ (CAC) function in *Morpho* (Schlager 2017). Consistent with the Gould-Mosimann framework for allometric accounting in geometric morphometrics (Klingenberg 2016), this method performs a multivariate regression of shape on centroid size to extract residual shape components (RSCs) that represent size-independent shape variation. We visualized allometric relationships by plotting centroid and other size variables against PC1 (FIG 3., S4 Figure, A-D). To validate the assumption that PC1 represented allometric shape change, we directly compared PC1 scores with CAC scores using a correlation test (S3 Figure).

#### [2.3f] Size-mediated and Size-independent Visualizations

We visualized patterns of cranial shape variation by calculating mean shapes from the Procrustes aligned coordinates for each habitat. We used the ‘findMeanSpec’ function in *geomorph* to identify the specimen in the dataset whose landmark coordinates were closest to the consensus mean configuration established by the GPA and used it as the reference specimen for all visualizations (FIG 4). A 3D mesh of the reference specimen was first warped to the overall mean and then plotted to the mean configurations for each group. The ‘meshDist’ function was used to create heatmaps with color representing mesh-to-mesh Euclidean distances of each group from the sample mean. To visualize patterns of shape variation between urban and rural environments in a size-free shape space, we back-projected the RSCs into Procrustes space and again calculated group means for the size-corrected data (FIG 4).

## 3. RESULTS

### [3.1] Body Size

#### [3.1a] Urban mice weigh less than rural mice

Habitat significantly contributed to variation in body weight analyses with [*P* = 0.007] and without [*P* = 0.015] body length as a covariate (S6 Table). Across both models, all other factors and interaction terms were not significant, except for body length in the model where it was a covariate [*P* < 0.001]. Emmeans (EMMs) comparisons of the covariate model revealed a difference in weight of 1.74g [*P* = 0.029] between urban (14.4g) and rural (16.1g) populations, suggesting that, even when accounting for body length, urban mice weigh ∼ 10.8% less on average than rural mice (FIG 2A). Here and for all subsequent linear models, assumptions of normality and homogeneity of variance were met (Shapiro-Wilk, Levene’s test *P* > 0.05).

**Figure 2.**
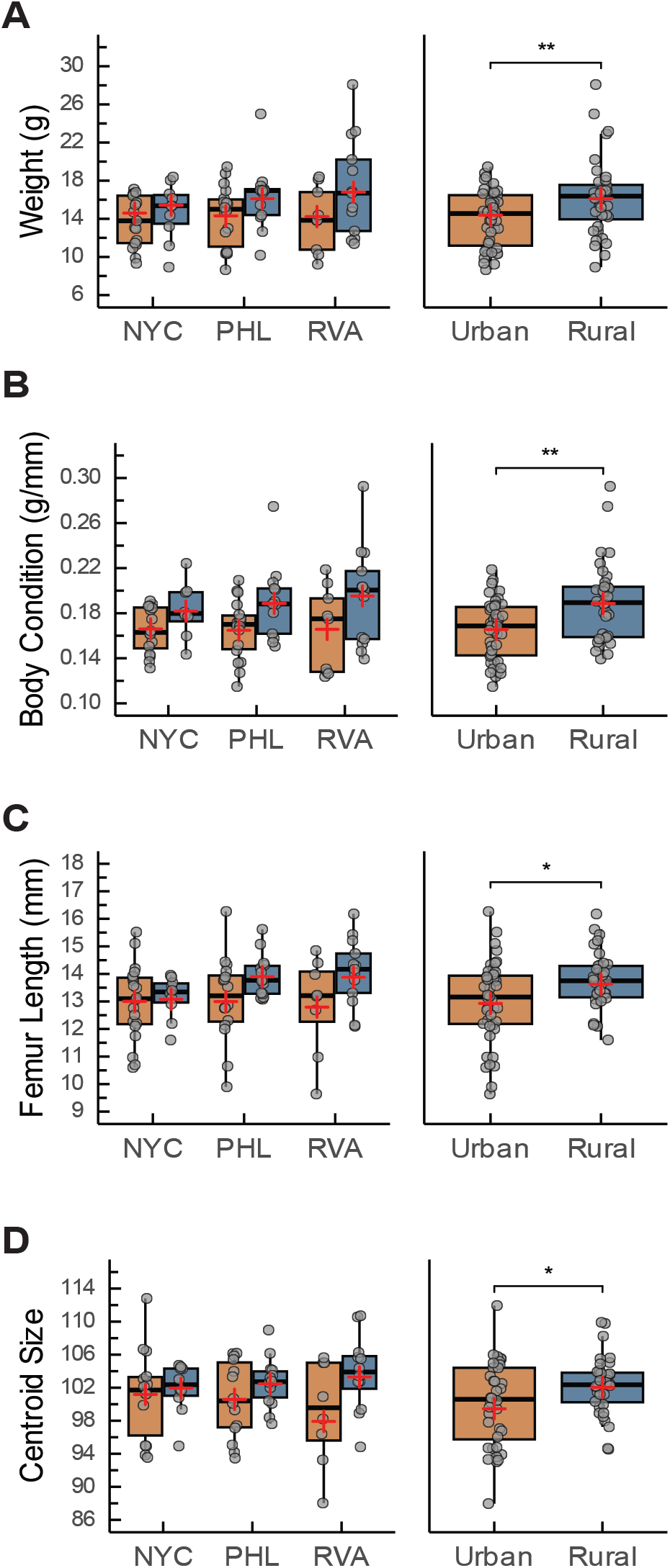
Urban mice are smaller than rural mice across several key measures. Boxplots are shown for (A) weight, body condition, (C) femur length, and (D) cranial centroid size within each habitat and metro region and aggregated over metro regions (^*^*P* < 0.05, ^**^*P* < 0.01). (**+**) symbols indicate the estimated marginal mean for each group when averaged over other factors as predicted by emmeans.

**Figure 3.**
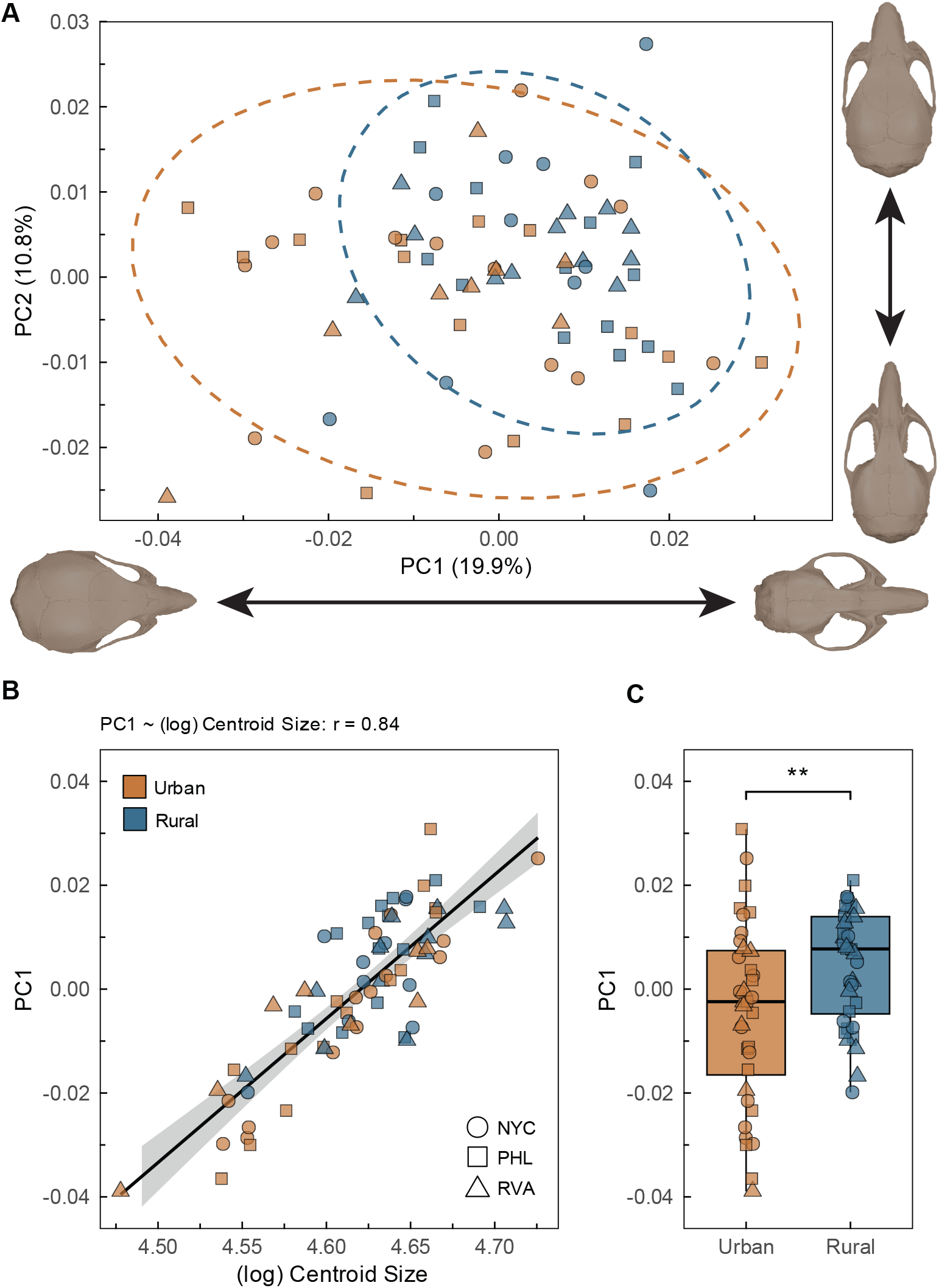
Allometry drives differences in cranial shape across habitat. (A) PCA plot of the first two principal components accounting for ∼ 30% of the overall variation in cranial shape. PC1 is the allometric shape axis. Exaggerated cranial warps are depicted to demonstrate shape changes associated with the extremes of each axis. (B) The strong positive linear relationship between (log) Centroid Size and PC1 indicates an allometric scaling effect. (C) PC1, which captures cranial shape differentiation driven by size, varies across habitat (^**^*P* < 0.01). Boxplots show the distribution of PC1 scores within urban and rural habitats.

**Figure 4.**
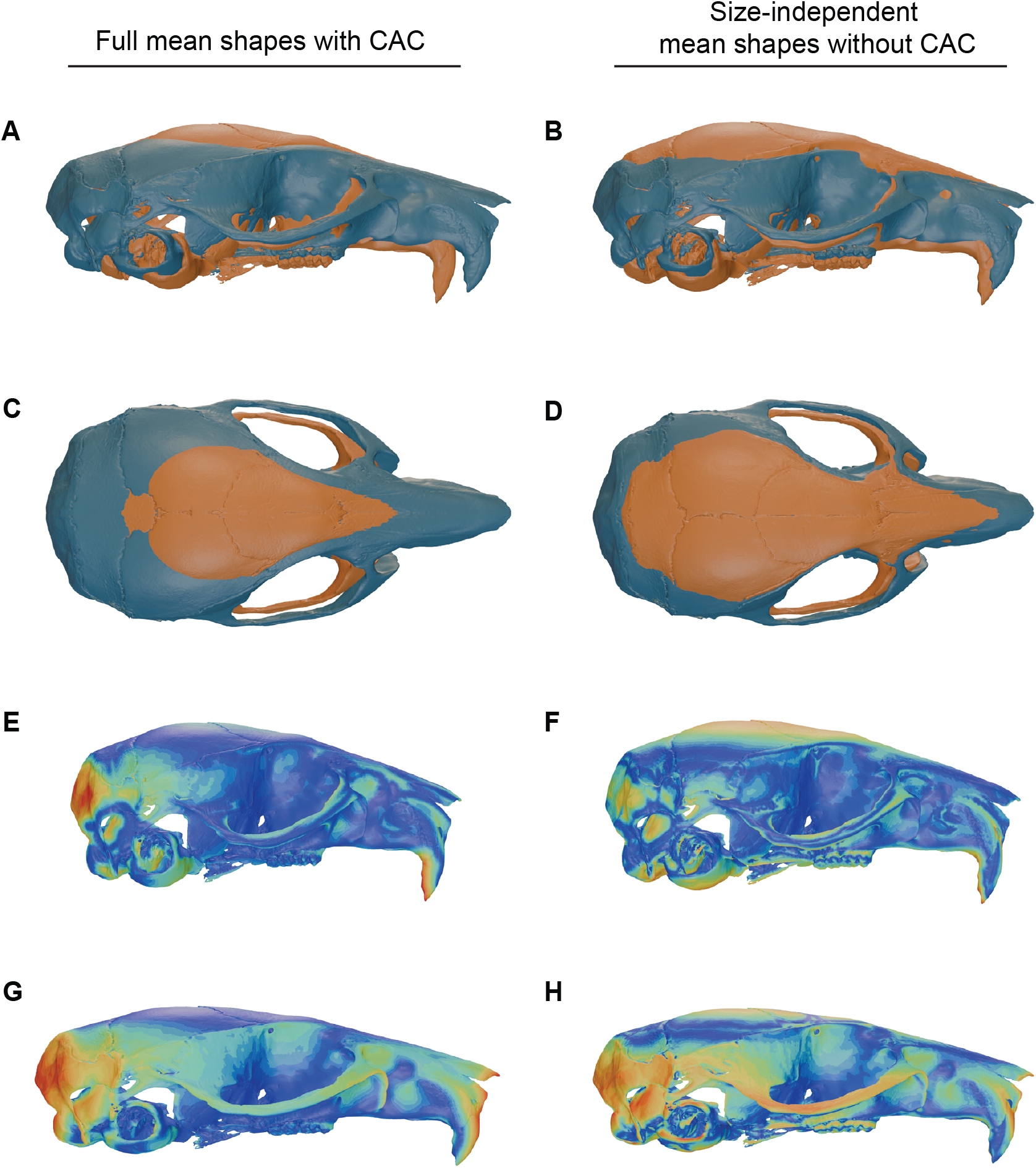
Urban and rural mice differ in cranial shape with variation both mediated by size (left column) and independent of size (right column). Mean shapes are shown with [A,C,E,G] and without [B,D,F,H] the common allometric component (CAC) for the habitat variable. Panels A-D show overlapping mean shapes in both lateral (A,B) and dorsal (C,D) views for comparison. Colors in panels [A, B, C, D] differentiate habitat (orange = urban, blue = rural). Panels E-H show urban (E,F) and rural (G,H) heatmaps, with warmer colors indicating regions of greater distance from the mean, and cooler colors indicating similarity. Shape differences are magnified 20x. Skulls in panel (C) were symmetrized for visual clarity. Unedited meshes are provided in Supplementary Figure S5.

We also considered body condition (body weight divided by length). Results of this analysis were similar, with habitat contributing significantly to variation [*P* = 0.003] but not additional factors or interaction terms. Contrasts implemented in emmeans predicted a 0.023g/mm difference in EMMs of urban and rural populations [*P* = 0.015] with body condition of urban mice estimated as ∼12.2% lower than their rural conspecifics. Overall, these results suggest that urban mice weigh less and are lighter per unit length compared to rural mice (FIG 2B).

#### [3.1b] Urban mice are smaller than rural mice

We analyzed body length, femur length, and femoral mediolateral width to assess linear size variation. First, we considered body length and found that no factors or interactions including habitat [*P* = 0.114] contributed significantly to variation. However, this measure can be noisy because it is typically calculated by measuring the total length of the specimen (from tip to tail) and subtracting a second measure for the tail (Jewell and Fullagar 1966; Di Masso et al. 1991). As a result, there are two opportunities for error. In addition, it is collected in the field using a ruler with lower precision than digital calipers. Therefore, we also analyzed femur length, which is collected in the lab after specimen preparation as a proxy for overall body length (Melin et al. 2005). Body length and femur length were significantly correlated [r = 0.71, *P* < 0.001]. We found that habitat, but no other factors or interactions, contributed significantly to variation in femur length [*P* = 0.015], indicating a reduction in overall body size in the urban populations (FIG 2C).

The effect of habitat on femoral mediolateral width [*P* = 0.854] was not significant in a model tested without femur length included as a covariate, though sex was a significant predictor [*P* = 0.041). However, in the covariate model, both habitat [*P* = 0.019] and sex [*P* < 0.001] were significant. Femur length also significantly predicted femoral mediolateral width [*P* < 0.001]. With results averaged over the other factors, emmeans contrasts revealed that rural mice (estimate: -0.055 mm, *P* = 0.125) and females (estimate: -0.103 mm, *P* = 0.004) had more gracile (i.e., thinner) femora than urban mice and males, respectively. Overall, these findings provide evidence for linear size differences between urban and rural mice, with urban mice on average having shorter and more robust femora than their rural counterparts, an indication of reduced body size.

### [3.2] Cranial Size and Shape

#### [3.2a] Consistent with body size, urban mice have smaller crania

Habitat was a significant predictor of centroid size [*P* = 0.035], but not when tested in a model with body length included as a covariate [*P* = 0.178]. In the covariate model, body length significantly predicted centroid size [*P* < 0.001]. No other factors or interaction terms were significant. Emmeans comparisons revealed that, consistent with linear size metrics, urban mice (estimate: 99.9) have lower estimated marginal mean centroid sizes (*P* = 0.043) than their rural counterparts (estimate: 102.6) (FIG 2D). Thus, cranial size differs among habitats, but that difference is explained by differences in body size.

#### [3.2b] Cranial shape variation is largely explained by allometry

First, we tested whether habitat, metro region, sex or size contributed to variation in cranial shape. Procrustes ANOVA with the aligned coordinates as a response variable revealed that centroid size was the top predictor of cranial shape in the full model [*Rsq* = 13.1%, *P* = 0.001]. With centroid included as a covariate, habitat [*Rsq* = 2.3%, *P* = 0.02] and metro region [*Rsq* 3.8%, *P* = 0.009] also contributed significantly to variation. Sex and associated interaction terms were not significant (S7 Table).

We also considered a similar model via MANCOVA using the PCA to summarize variation in shape. Parallel analysis of the PCs indicated that the first 9 PCs captured meaningful shape variation (S5 Table, S2 Figure) and these PCs were retained (PC9 proportion of variance = 2.8%; cumulative proportion of variance = 66.4%, S3 & S4 Tables). The first three PCs (PC1-3) accounted for 19.9%, 10.8% and 10.7% of the variation, respectively (S4 Table). MANCOVA results were concordant with the findings from the Procrustes ANOVA, with centroid size, habitat, and metro region [all *P* < 0.001] identified as significant predictors (S8 Table).

The significant contribution of centroid size to shape variation suggested that allometric scaling effects may be important in this system. We tested a reduced allometric model and found that the centroid:habitat interaction term was not significant (S3 Figure), indicating that urban and rural mice occupy a shared allometric slope. Therefore, we visualized shared shape∼size covariation by plotting centroid on the first PC axis (FIG 3B). We verified that PC1 captured the size-related shape variation by directly comparing CAC scores with PC1 scores (Pearson’s r = 0.99, S3 Figure). This strong association suggests that the primary axis of shape variation adequately captures the allometric signal, and that PC1 can be effectively treated as the allometry axis.

The positive linear relationship between centroid and PC1 indicates that skull shape changes systematically with size, with larger crania associated with positive PC1 scores and smaller crania associated with negative PC1 scores (FIG 3B) Differential placement along the shared vector suggests that urban and rural mice occupy distinct regions of cranial morphospace according to their size (Welch’s two-sample t-test [*P* = 0.009], FIG 3C) We plotted other body size metrics against PC1 to visualize their association with cranial shape (S4 Figure, A-D). These variables also showed a positive linear relationship with PC1, suggesting that the primary axis of cranial shape variation reflects general body size allometry and not scaling patterns specific to the size of the skull. Shape differences along PC1 are associated with changes in the length of the rostrum, width and angle of the zygomatic region, and globularity of the brain case (FIG 3A).

Visual comparison of habitat mean shapes suggests that urban mice have dorsoventrally expanded and craniocaudally compressed skulls—that is that urban mice have taller skulls that are shorter along the anterior-posterior axis (FIG 4, A,C,E,G). In comparison, rural crania are longer along the craniocaudal axis, resulting in a flatter and wider appearance. Additionally, restriction along the craniocaudal axis in the urban population suggests shorter molar tooth rows, and a relatively shorter distance between the tooth rows and the most anterior aspect of the rostrum. Conversely, rural crania show an increase in this distance, resulting in longer rostra when compared to the urban mean.

#### [3.2c] Urban and rural mice differ in cranial shape after accounting for size

The significant effect of habitat in both the full and reduced allometry models indicates a shift of the slope resulting in different intercepts between the populations (S3 Figure). To interpret size-independent cranial shape differences associated with the intercept shift between habitats, we extracted the CAC from the shape data and again generated an overall mean configuration from the global population. This configuration was then warped to the population means to visualize habitat-specific patterns of skull shape (FIG 4, B,D,F,H). Size-independent mean shape comparisons show a reduction in the magnitude of difference between urban and rural shapes along the craniocaudal axis. Additionally, urban crania appear even more vertically expanded, with dorsal and ventral surfaces visible beyond the rural mesh (FIG 4, B & D). The vertically taller and deeper skull also results in a comparatively greater slope extending from the cranial vault to the rostrum in the urban population.

### 3.3 Cranial Endocast Volume

#### [3.3a] Variation in cranial endocast volume is driven by body size

Habitat did not significantly contribute to variation in endocast volume in models with [*P* = 0.849] or without [*P* = 0.078] cranial centroid size included as a covariate. Interactions between metro region and habitat [*P* = 0.047] and sex and habitat [*P* = 0.032] were marginally significant only in the model without centroid size as a covariate. However, centroid size was a significant predictor when included as a covariate [*P* < 0.001; S6 Table] and volume and cranial centroid size were highly correlated (S4 Figure). Overall, results suggest that variation in cranial endocast volume are driven by variation in body size. Differences in cranial volume between habitats are consistent with what would be predicted given differences in size. Therefore, there is no evidence for house mice having relatively greater endocranial volume in urban environments.

## 4. DISCUSSION

We investigated morphological variation between urban and rural populations of *M. m domesticus* across three U.S. cities and found significant variation in multiple traits. Urban mice weigh less overall and per unit length compart to their rural conspecifics. In addition, urban mice have shorter femora (a proxy for body length) and smaller cranial centroid sizes (a proxy for skull size). Together, these results indicate that urban mice are smaller bodied overall and also weigh less for their body size. Our analysis of cranial shape revealed strong covariance between size and shape, with centroid size accounting for ∼13% of the variation in the permutation model. Metro region (3.8%) and habitat (2.3%) accounted for less variation but still emerged as significant predictors. These findings point to a clear signal of allometry, with most of the cranial shape variation attributable to size, but some allometry-free variation attributable directly to habitat and metro region. Importantly, however, urban and rural mice differ in size, meaning that the size-related shape variation itself can be interpreted indirectly as a habitat effect.

### [4.1] Ecological drivers of body size differentiation and variation in taxonomic responses to urbanization

Here, we found that cities were associated with reduced body size and condition. Meta-analyses of body size variation in response to urbanization suggest that overall, rodents tend to be larger in urban environments (Santini et al. 2019; Hantak et al. 2021). Nevertheless, results from studies of individual rodent taxa are mixed. Some, like ours, have found that body size metrics are negatively correlated with urbanization class or relevant proxies (i.e. human population density), including body length [*Peromyscus maniculatus* (Guralnick et al. 2020); *Peromyscus leucopus & Tamias striatus* (Lopez 2024)], body mass [*Mus musculoides* (Ofori et al. 2024); *Sciurus vulgaris* (Wist et al. 2022)], and body condition [*Mastomys natlensis, Mus musculoides*, & *Tatea kempi* (Ofori et al. 2024); *Chaetodipus* spp., *Perognathus* spp., & *Dipodomys merriami* (Bateman et al. 2023); *Sciurus vulgaris* (Wist et al. 2022)]. On the other hand, studies have identified increases in aspects of body size associated with urbanization [e.g. *Apodemus agrarius* (Liro 1985; Pieniążek et al. 2017); *Sciurus carolinensis* (Tranquillo et al. 2024)].

The lack of consistency across studies suggests that the factors driving variation in body size are likely diverse and the effects of those factors are also likely mediated by diverse life histories within the very speciose Rodentia. Body size is an important functional trait tied to multiple aspects of a species’ ecology and life history, including metabolic rate, thermoregulatory ability, diet, predation, vagility, fecundity, among others (Kleiber 1932; Blueweiss et al. 1978; Western 1979; Cohen et al. 1993)). There are many factors, both abiotic and biotic, associated with urbanization that may be predicted to impact body size. For example, physiological constraints imposed by body size may create energetic challenges in response to the thermal environment. These thermoregulatory and energetic dynamics are thought to underlie clinal patterns of body size variation where endotherms are larger in colder environments and smaller in warmer ones. This ecogeographic pattern, formally known as Bergmann’s rule (Bergmann 1847), has been repeatedly established in mammalian taxa (Meiri and Dayan 2003; Blackburn and Hawkins 2004), including in house mice (Lynch 1992; Phifer-Rixey et al. 2018). The application of Bergmann’s rule in the context of urban heat islands (UHI) predicts smaller body sizes in comparatively warmer urban environments and has been a recent focus in the urban ecology literature (Merckx et al. 2018; Hantak et al. 2021; Bateman et al. 2023).

However, there are many other factors to consider, including diet, predation, and dispersal/range size, that may contribute to body size variation across urban and non-urban habitats. Dietary shifts between habitats can occur as urban populations exploit more anthropogenic resources, and the quantity and quality of resources available to urban populations can either increase or decrease body size (McCleery 2015; Martin and Sheridan 2022). Stable isotope analyses in these *M. m. domesticus* populations has uncovered diets higher in animal protein in cities along with associated changes in gut microbial composition (Giancarli et al. 2026). Rural mice are expected to be primarily dependent on plant-based animal feed while urban mice likely have greater access to human-derived food sources and protein-based companion animal feed. Urbanization can also alter predation and has been shown to result in an increase in predator density, but an overall decrease in predation pressure; “the predation paradox” (Fischer et al. 2012). This release from predation may be predicted to result in larger body size. On the other hand, in commensal species, this may not be the case. Commensals like house mice are likely more susceptible to direct encounters with humans and domesticated animal predators which can result in high mortality (Woods et al. 2003). Therefore, a reduction in body size (crypsis) may be a strategy to avoid detection and ultimately predation. Finally, dispersal ability and range size are positively correlated with body size (Lindstedt et al. 1986; Sutherland et al. 2000; Santini et al. 2013; Martin and Sheridan 2022). In urban habitats, species with foraging habits or reproductive strategies that require large ranges and/or dispersal may experience pressure for enhanced vagility, especially if habitat patches are unevenly distributed or fragmented across the landscape. This hypothesis likely has little relevance for house mice which are not expected to disperse over large distances with human assistance (Mikesic and Drickamer 1992; Pocock et al. 2005), but other rodents have relatively large home ranges and can traverse large distances despite structural barriers in urban habitats (Tolkachev 2016; Dammhahn et al. 2020). It is clear that at finer taxonomic scales, the effect of urbanization on body size can be heterogeneous (Callaghan et al. 2025). In order to achieve consensus, more studies like ours that investigate new species and utilize paired replicate designs to identify convergent patterns will be needed to uncover the factors that drive body size variation.

### [4.2] Functional consequences of cranial shape associated with diet

We uncovered size-mediated and size-independent patterns of cranial shape variation that may be associated with diet since functional demands for biting and mastication can be associated with cranial shape (Marcy et al. 2024; Mitchell et al. 2024). Generally, flatter crania with longer rostra as observed in the rural phenotype are associated with reduced bite force capacity and lower mechanical efficiency (Mitchell et al. 2024; Chandler et al. 2025). Further, a dorsoventrally deeper skull, which is a substantial morphological trend observed in both the size-mediated and size-independent components of shape for urban mice (FIG 4), provides increased reinforcement against stresses associated with jaw adduction (Mitchell et al. 2024).

Additionally, rural mice appear to have longer molar tooth rows than urban mice (FIG 4A). Longer molar tooth rows are associated with low-quality diets and the consumption of food resources that require more chewing (Yu et al. 2017). These results are consistent with dietary differences among habitats in our populations (Giancarli et al. 2026). Interestingly, this pattern of shorter molar tooth rows in association with urban habitats has been uncovered in the white-footed mouse [*Peromyscus leucopus* (Yu et al. 2017)] and brown rat [*Rattus norvegicus* (Puckett et al. 2020)] in New York City.

### [4.3] Variation in cranial endocast volume is consistent with differences in body size

Increased cranial capacity has been posited as a response to urban structural complexity in rodents (Snell-Rood and Wick 2013). In our study, habitat did not directly contribute to variation in endocast volume. Instead, endocast volume was significantly predicted by centroid size, a proxy for body size. Interestingly, our data suggest that in urban *M. m. domesticus*, endocast volume is smaller due to reduced body size—an indirect effect of habitat. It is perhaps not surprising that our results do not support the hypothesis of increased cranial capacity in urban contexts. Evidence for the hypothesis in rodents has been limited to date (Snell-Rood and Wick 2013; DePasquale et al. 2020) and, overall, studies suggest that responses to urbanization for this trait are variable (Snell-Rood and Wick 2013; Puckett et al. 2020). Variation in response may be driven by diverse life histories and evolutionary histories in rodents (Kay and Hoekstra 2008; Fabre et al. 2012). Notably, in their original paper, Snell-Rood and Wick identified only two out of ten species as having relatively larger cranial capacities in cities, and for neither of those did they also find body size differences among habitats. The authors also suggest that trends toward increased cranial capacity may be most prevalent in the early stages of colonization to an urban habitat (Snell-Rood and Wick 2013). House mice are human commensals that have occupied complex anthropogenic environments for thousands of years, and more recently in burgeoning pre-urban and urban settlements in both the western and eastern hemispheres (Boursot et al. 1993).

### [4.4] Allometric patterns in cranial morphology

Our results uncovered a strong allometric signal of cranial shape differentiation mediated by body size. Allometry is the covariation of a trait with body size which occurs over the course of an individual’s development (*ontogenetic*) but also potentially between populations of the same species at a similar developmental stage (*static*) (Gould 1971; Cheverud 1982; Mitchell et al. 2024). The observed size-mediated shape variation (FIG 3, FIG 4) indicates that urban mice occupy a distinct region of cranial morphospace along a shared allometric trajectory, reflecting differences in body size between habitats. This pattern implies that divergence in cranial shape between habitats is partly structured by size-related constraints. Importantly, in the context of the results presented in (3.1) that show significant signals of size differentiation between urban and rural populations, the size-mediated shape variation itself can be interpreted indirectly as a habitat effect. Because the mice sampled in this study were developmentally mature, we interpret these findings as a signal of static allometry between urban and rural habitats.

It is nominally possible that we have disproportionately sampled younger mice in urban environments. While accurately assessing age in wild mice is challenging, we used hindfoot length ≥ 18mm as a developmental indicator of maturity. This cut-off represents a conservative baseline within the reported adult range of 15-20mm for the species. While we cannot fully rule out the possibility that the results observed here reflect some amount of ontogenetic allometry, this explanation is unlikely to account for the consistent patterns observed across all three cities or for the differences in body condition.

With respect to cranial shape allometries, predictable and repeatable patterns have been identified in mammals. For example, placental mammals broadly follow patterns of Craniofacial Evolutionary Allometry (CREA), where larger species tend to have relatively longer faces and smaller brain cases (Radinsky 1985; Cardini and Polly 2013; Cardini 2019). While CREA has been established as a broad evolutionary pattern, less is known about its relevance on microevolutionary scales. Still, some comparative work has described intraspecific CREA (Schlis-Elias and Malaney 2022; Rhoda et al. 2023; Chandler et al. 2025), suggesting that the pattern is not a strictly macroevolutionary phenomenon. In our dataset, shape changes toward the positive end of PC1 are associated with an elongation of the face/rostrum and a relative constriction of the brain case. Mean shapes in FIG 4 also show this elongation along the craniocaudal axis in the larger-bodied rural mice and are consistent with the CREA pattern. The prevalence of size-related cranial shape variation in our dataset underscores the need for researchers interested in craniofacial phenotypes in the context of urbanization to consider body size as a potential confounding factor in their analyses.

## 5. CONCLUSION

Studies of morphological variation like ours are especially important for common urban taxa which are often considered pests and underrepresented in museum collections (Shultz et al. 2021; Winchell et al. 2022). Nevertheless, while phenotypic variation in wild house mice remains poorly characterized (but see Boell and Tautz 2011; Pallares et al. 2016; Parmenter et al. 2020; Ballinger and Nachman 2022; Alibert 2025), especially in cities, house mice have great potential as a vertebrate model of urban evolutionary ecology. The global distribution of house mice allows for the study of populations across a range of environmental contexts, facilitating investigation of how urbanization interacts with other factors (e.g. climate, latitude) and helping identify parallel, divergent, and idiosyncratic responses. In addition, because house mice are a genetic model system, there is a strong foundation for future genotype-phenotype and genotype-environment associations in the context of urban-associated morphological variation. For example, both body size and cranial shape are complex quantitative traits with a discrete genetic basis that has been extensively explored in both wild-derived and classical inbred strains (MacArthur 1949; Cheverud et al. 1996; Keightley et al. 1996; Kenney-Hunt et al. 2006; Wuschke et al. 2007; Chan et al. 2012; Maga et al. 2015; Pallares et al. 2015).

Our results provide strong evidence of urban-associated morphological differentiation and point to body size as a key trait mediating response to habitat. Studies, like ours, that comprehensively investigate morphology in a replicated paired urban-rural framework remain rare in vertebrates. Importantly, while the differences observed in this study could result from many factors (genetic variation, environmental variation, phenotypic plasticity, genotype-by-environment interactions, or a combination), the consistent signal across habitats makes it unlikely that patterns result from neutral processes like genetic drift. Moreover, whatever the mechanism, the consistent and meaningful differences in body size across habitat signal strong potential for impacts on fitness, motivating future studies.

## Funding

This work was supported by the National Science Foundation [grant number 2332998 to M.P.R.].

## Acknowledgements

We would like to thank Dr. Kristof Zyskowski and Piper Stepule of the Yale Peabody Museum for their efforts in specimen processing. Additionally, we thank the Penn Center for Musculoskeletal Disorders (PCMD) MicroCT Imaging Core [NIH/NIAMS P30 AR069619] for instrument access, and manager Dr. Wen Sang for assistance with the micro-CT workflow. We also thank lab members Dr. Adrienne Kasprowicz, Dr. René Clark, Dr. Sam Giancarli, Nina Mallalieu, Zade Alafranji, and Jessica Bonnen for their feedback and support, as well as undergraduates Julian Atendido and Angele Oye-Mba for their help in preliminary analyses. Finally, we are especially grateful to the many community participants who have allowed us into their homes and farms to trap the mice used in this project.

## Data Availability

Data underlying this article are made available in its online supplementary material as well as on Dryad [in progress]. Digital 3D cranial renderings for relevant specimens have been uploaded to MorphoSource [in progress]. Code used for analyses and figures can be viewed at https://github.com/PhiferrixeyLab/Kupchella_2026.

## Notes

### Competing Interest Statement

The authors have declared no competing interest.

