## Supplementary Figures for "Body size and cranial shape differentiation in urban and rural house mice (*Mus musculus domesticus*)"

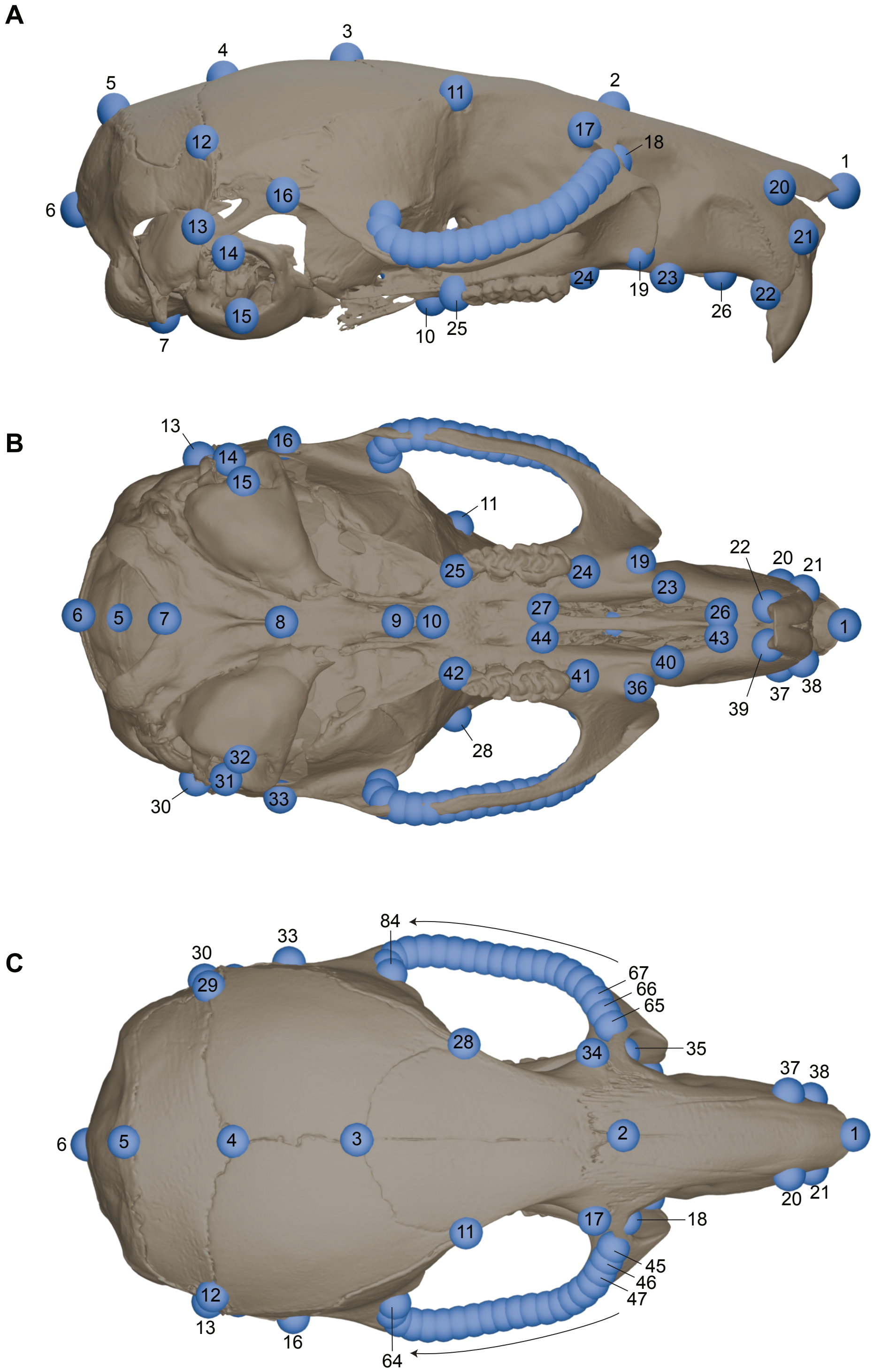


**Supplementary Figure S1.** *Landmark Map.* **A)** Right lateral, **B)** ventral, and **C)** dorsal views of a map of the 84 landmarks used in the cranial shape analysis.

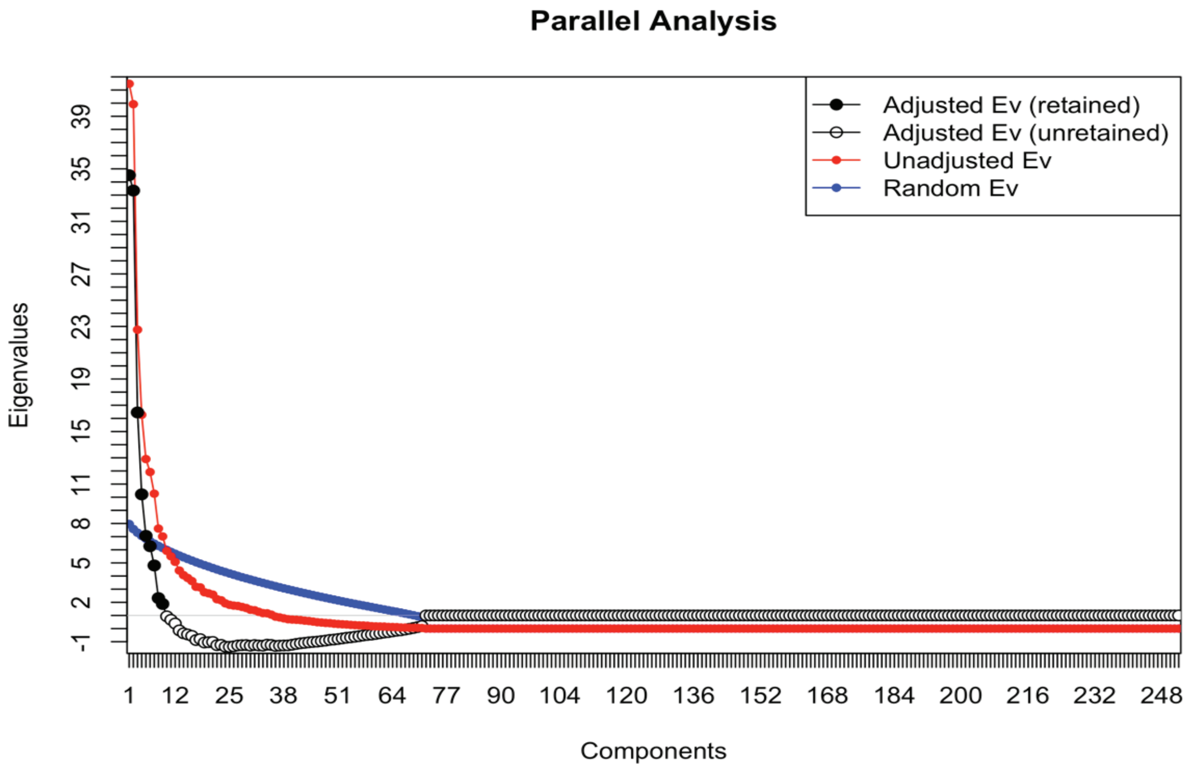


**Supplementary Figure S2.** *Parallel Analysis Scree Plot*. Nine PCs (black dots), whose adjusted eigenvalues were >1 in a Horn's Parallel analysis in the *paran* package, were retained.

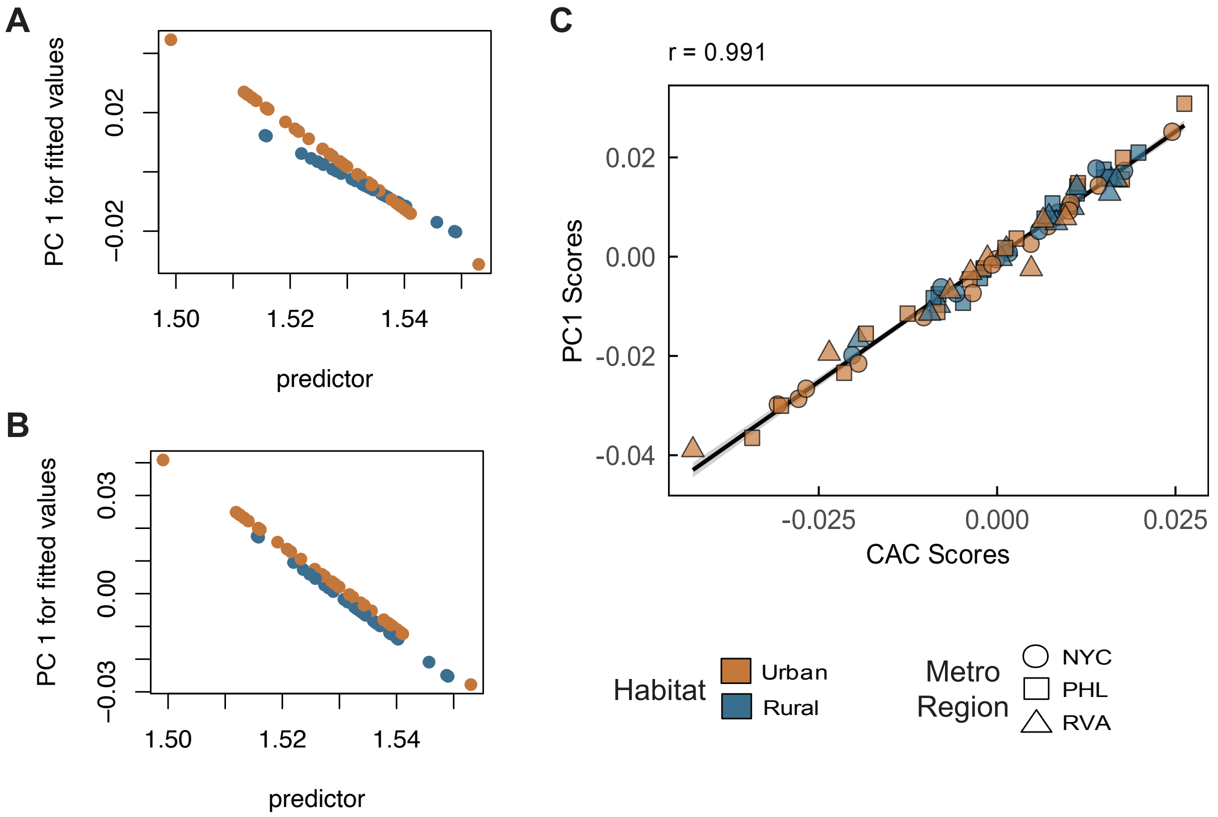


**Supplementary Figure S3**. *Reduced allometry models and CAC/PC1 correlation.* (A) The log_centroid:habitat interaction term was not significant ($P = 0.761$) in the plotAllometry model with habitat included as an interaction term (shape ~ size * habitat), providing evidence that urban and rural mice share an allometric slope. Therefore, we used (B) the simpler plotAllometry model with habitat included as an additive term (shape ~ size + habitat), where slopes are presumed the same and habitat is significant ($P = 0.017$), showing different intercepts. (C) There is a strong positive linear association between PC1 and CAC scores (Pearson's r = 0.991), suggesting that PC1 captures the allometric relationship between size and cranial shape variation.

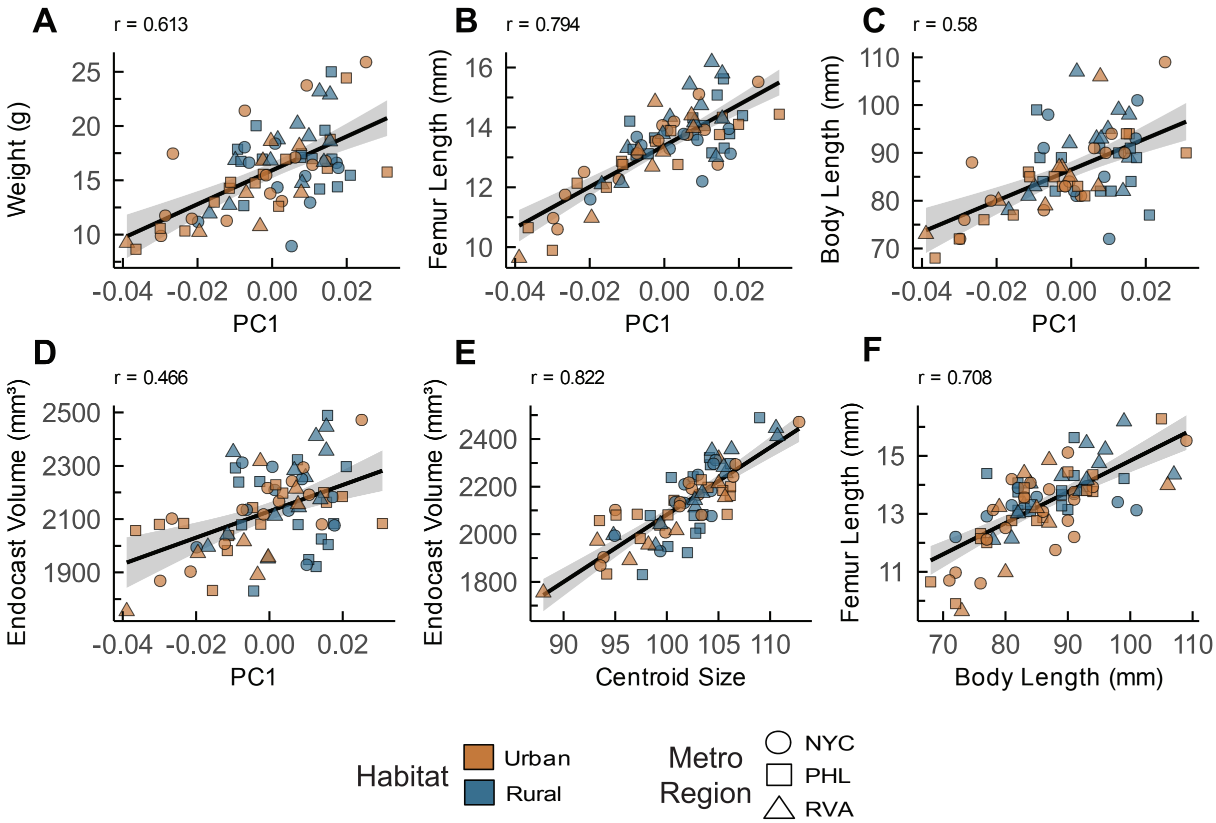


**Supplementary Figure S4.** *Allometric correlations.* Plots showing the relationship between size and shape variables. Panels (A-D) show the relationship between cranial shape PC1 and (A) body weight, (B) femur length, (C) body length, and (D) cranial endocast volume. Panels E and F show the positive linear association between (E) cranial endocast volume and cranial centroid size, and (F) femur length and body length. Pearson's correlation test was performed for each comparison. In all cases, $P < 0.001$ and each plot is labeled with the Pearson correlation coefficient.

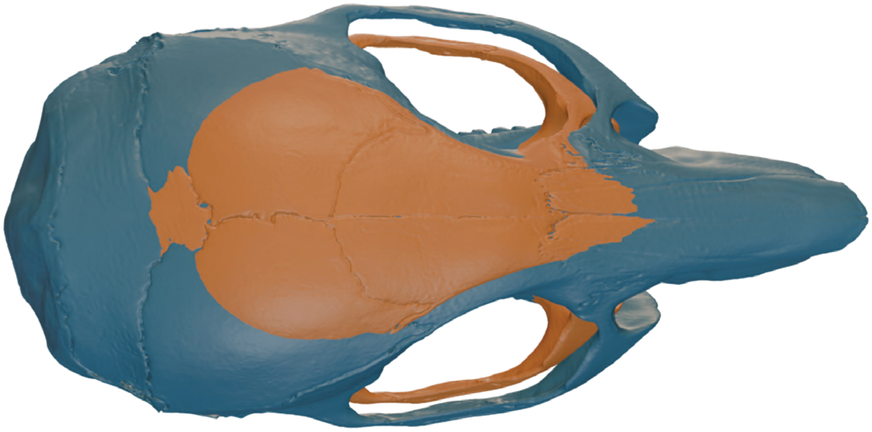

**Supplementary Figure S5.** *Unedited full mean shapes with CAC, dorsal view.* A dorsal view of the full mean shapes from Figure 4C retaining minor asymmetry. Skulls in Figure 4C were symmetrized for visual clarity.
